# Coexistence of multiple endemic and pandemic lineages of the rice blast pathogen

**DOI:** 10.1101/179895

**Authors:** Pierre Gladieux, Sébastien Ravel, Adrien Rieux, Sandrine Cros-Arteil, Henri Adreit, Joëlle Milazzo, Maud Thierry, Elisabeth Fournier, Ryohei Terauchi, Didier Tharreau

## Abstract

The rice blast fungus *Magnaporthe oryzae* (syn. *Pyricularia oryzae*) is both a threat to global food security and a model for plant pathology. Molecular pathologists need an accurate understanding of the origins and line of descent of *M. oryzae* populations, to identify the genetic and functional bases of pathogen adaptation, and to guide the development of more effective control strategies. We used a whole-genome sequence analysis of samples from different times and places to infer details about the genetic makeup of *M. oryzae* from a global collection of isolates. Analyses of population structure identified six lineages within *M. oryzae*, including two pandemic on japonica and indica rice, respectively, and four lineages with more restricted distributions. Tip-dating calibration indicated that *M. oryzae* lineages separated about a millenium ago, long after the initial domestication of rice. The major lineage endemic to continental Southeast Asia displayed signatures of sexual recombination and evidence of DNA acquisition from multiple lineages. Tests for weak natural selection revealed that the pandemic spread of clonal lineages entailed an evolutionary ‘cost’, in terms of the accumulation of deleterious mutations. Our findings reveal the coexistence of multiple endemic and pandemic lineages with contrasting population and genetic characteristics within a widely distributed pathogen.

**Importance:** The rice blast fungus *Magnaporthe oryzae* (syn. *Pyricularia oryzae*) is a textbook example of a rapidly adapting pathogen, and is responsible for one of the most damaging diseases of rice. Improvements in our understanding of *Magnaporthe oryzae* diversity and evolution are required, to guide the development of more effective control strategies. We used genome sequencing data for samples from around the world to infer the evolutionary history of *M. oryzae.* We found that *M. oryzae* diversified about a thousand years ago ago, separating into six main lineages: two pandemic on japonica and indica rice, respectively, and four with more restricted distributions. We also found that a lineage endemic to continental Southeast Asia displayed signatures of sexual recombination and the acquisition of genetic material from multiple lineages. This work provides a population-level genomic framework for defining molecular markers for the control of rice blast and investigations of the molecular basis of differences in pathogenicity between *M. oryzae* lineages.

## Introduction

Fungal plant pathogens provide many examples of geographically widespread, often clonal, lineages capable of adapting rapidly to anthropogenic changes, such as the use of new fungicides or resistant varieties, despite extremely low levels of population genetic diversity [1, 2]. An accurate characterization of the population biology and evolutionary history of these organisms is crucial, to understand the factors underlying their emergence and spread, and to provide new, powerful and enduring solutions to control these factors. A knowledge of the origins and lines of descent connecting extant pathogen population provides insight into the pace and mode of disease emergence and subsequent dispersal [2, 3]. By inferring the history and structure of pathogen populations, we can also identify disease reservoirs and improve our understanding of the transmissibility and longevity of populations [4, 5]. Finally, quantification of the amount and distribution of genetic variation across space and time provides a population-level genomic framework for defining molecular markers for pathogen control and for investigations of the molecular basis of differences in phenotype and fitness between divergent pathogen lineages.

Rice blast is one of the most damaging rice disease worldwide [6-8]. It is caused by the ascomycete fungus *Magnaporthe oryzae* (syn. *Pyricularia oryzae*), which has become a model for plant pathology in parallel with the development of rice as a model crop species [7, 9-11]. The rice-infecting lineage of *M. oryzae* coexists with multiple host-specialized and genetically divergent lineages infecting other cereals and grasses [12-14]. The lineage infecting foxtail millet (*Setaria italica*, referred to hereafter as “Setaria”) is the closest relative of the rice-infecting lineage and rice blast was, thus, thought to have emerged following a host shift from Setaria about 2500 to 7500 years ago [15], at a time when Setaria was the preferred staple in East Asia [16, 17]. *Magnaporthe oryzae* infects the two major subspecies of rice, *Oryza sativa* ssp. *indica* and *japonica* (referred to hereafter as “indica” and “japonica”, respectively). Population genomics studies have provided support for a model in which *de novo* domestication occurred only once, to generate the japonica lineage, which subsequently diverged into temperate and tropical japonica, with introgressive hybridization from japonica leading to domesticated indica [18-20]. Using microsatellite markers, Saleh et al. [21] identified multiple endemic and pandemic genetic pools of rice-infecting strains, but were unable to resolve the evolutionary relationships between them. Rice blast has proved able to adapt rapidly to varietal resistance, and is thus a dynamic threat to such resistance in rice agrosystems [22]. This ability to adapt is surprising given the low level of diversity in *M. oryzae* and its infertility or asexual mode of reproduction in most rice-growing areas [22, 23]. This pathogen is, thus, particularly exposed to the “cost of pestification” (by analogy to the cost of domestication [24-27]), according to which, the combination of a small effective population size, strong selection for pestification genes and a lack of recombination lead to the accumulation of deleterious mutations [28]. Potential limitations to adaptation could be counterbalanced by boom-and-bust cycles in *M. oryzae*, with adaptation occurring during the boom phases, when short-term effective population size is large [2, 29]. Adaptive mutations may also be introduced by cryptic genetic exchanges with conspecifics or heterospecifics [30-33], but these mechanisms remain to be investigated in natural populations of *M. oryzae* [34]. An accurate understanding of the population genetics of successful clonal fungal pathogens, such as *M. oryzae*, can provide important insight into the genomic and eco-evolutionary processes underlying pathogen emergence and adaptation to anthropogenic changes.

We used pathogenicity data and whole-genome resequencing data for *M. oryzae* samples distributed over time and space, to address the following questions: What population structure does *M. oryzae* display? Does this species consist of relatively ancient or recent clonal lineages? What is history of temperate japonica, tropical japonica and indica japonica rice colonization by *M. oryzae?* Do *M. oryzae* lineages display differences in pathogenicity toward rice subspecies? Can we identify genetic exchanges between rice-infecting lineages and which genomic regions have been exchanged? Is there evidence for a cost of pestification in terms of the accumulation of deleterious mutations?

## Results

### Genome sequencing and SNP calling

We elucidated the emergence, diversification and spread of *M. oryzae* in rice agrosystems, by studying genome-wide variation across geographically widespread samples. We used 25 and 18 genomes sequenced by Illumina single-end and paired-end read technologies, respectively, with seven published genome sequences obtained with Solexa and mate-pair Titanium methods [10, 12]. We thus had a total of 50 genomes available for analysis (Table S1). Forty-five of the isolates concerned originated from cultivated rice (*Oryza sativa*), four from cultivated barley (*Hordeum vulgare*), and one from foxtail millet (*Setaria italica).* The sample set included multiple samples from geographically separated areas (North and South America, South, Southeast and East Asia, sub-Saharan Africa, Europe and the Mediterranean area), the reference laboratory strain 70-15 and its parent GY11 from French Guiana. Nine samples were collected from tropical japonica rice, seven from temperate japonica, fifteen from indica, and three from hybrid elite varieties. Sequencing reads were mapped onto the 41.1 Mb reference genome 70-15. Mean sequencing depth ranged from 5x to 64x for genomes sequenced with single-end reads, and from 5x to 10x for genomes sequenced with paired-end reads (Table S1). SNP calling identified 182,804 biallelic single-nucleotide polymorphisms (SNPs) distributed over seven chromosomes. The dataset consisted of 95,925 SNPs, excluding the Setaria-infecting lineage, 61,765 of which had less than 30% missing data, and 16,370 of which had no missing data.

### Population subdivision, genealogical relationships and levels of genetic variation

We used a multivariate analysis of population subdivision method, rather than model-based clustering algorithms, because multivariate methods require no assumptions about outcrossing, random mating or linkage equilibrium within clusters, and previous studies have shown that, in many populations, *M. oryzae* has lost its sexual recombination capacity ([21-23] and references therein). We used a discriminant analysis of principal components to determine the number of lineages represented in our dataset. Progressively increasing the number of clusters *K* from two to five identified the four lineages previously described by Saleh et al. in Asia [21] on the basis of microsatellite data, and a cluster of three strains collected from the Yunnan and Hunan Provinces of China (Figure 1). Further increases in *K* led to the subdivision of this Yunnan/Hunan cluster. Barley-infecting isolates clustered within rice-infecting lineage 1, confirming previous phylogenetic studies [12, 13]. Barley is “universally susceptible” to rice-infecting isolates, at least in laboratory conditions. However, the barley isolates included in this study were collected in Thailand, and no major blast epidemic has since been reported on this host in this area, indicating that barley is a minor host for rice-infecting populations.

**Figure 1.**
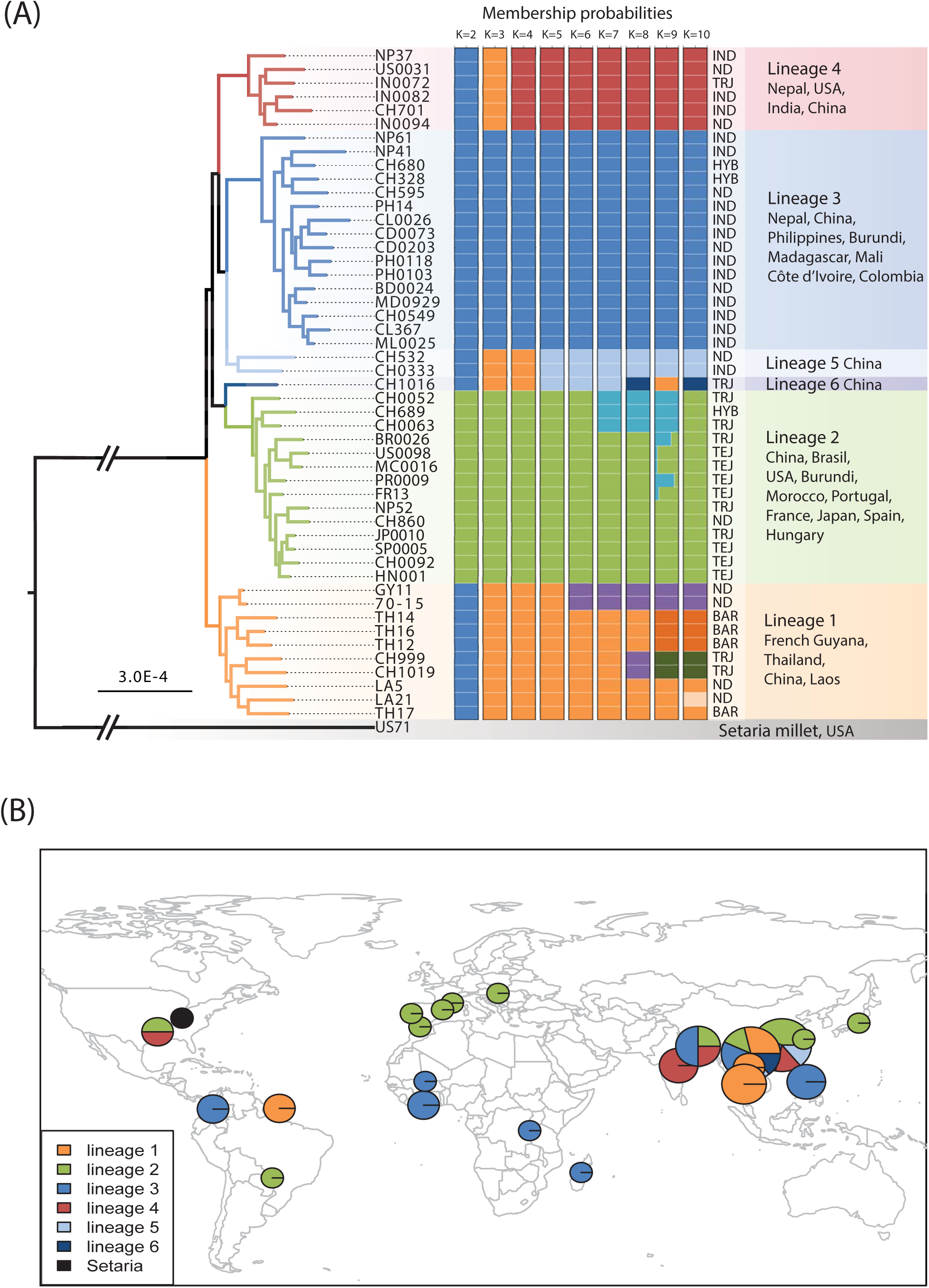
Population subdivision in the sample set analyzed. (A) Total-evidence maximum likelihood genome genealogy and discriminant analysis of principal components, (B) geographic distribution of the six lineages identified, based on results presented in (A). In panel A, all nodes had more than 95% bootstrap support (100 resamplings), except for the node carrying isolates BR0026, US0098, PR0009 and MC0016 (support: 72%). On the barplot, each isolate is represented by a thick horizontal line divided into *K* segments indicating the isolate’s estimated probability of belonging to the *K* assumed clusters. In panel B, diameters are proportional to the number of isolates collected per site (the smallest diameter represents 1 isolate). TRJ, tropical japonica; TEJ, temperate japonica; IND, indica; HYB, hybrid; BAR, barley; ND, no data.

We investigated whether the clusters observed at *K*>4 in the DAPC represented new independent lineages or subdivisions of the main clusters, by using RAxML to infer a genome genealogy [35]. We based the analysis on a dataset combining the full set of SNPs and monomorphic sites, rather than just SNPs, to increase topological and branch length accuracy [36]. The total-evidence genealogy revealed the existence of four lineages, corresponding to lineages 1 to 4 described by Saleh et al. [21], and two new lineages (named lineages 5 and 6) corresponding to the three-individual cluster observed at *K*=5 in the DAPC (Figure 1). With the 41 Mb dataset, including missing data, the most basal divergence within the rice-infecting lineage was that between lineage 1 and the other five lineages (Figure 1). If positions with missing data were excluded (15 Mb), the most basal divergence was that between a group composed of lineages 1, 2 and 6, and a group composed of lineages 3, 4 and 5 (not shown).

Absolute divergence (dxy) between pairs of lineages was of the order of 10^-4^ differences per base pair and was highest in comparisons with lineage 6 (Table S2). Nucleotide diversity within lineages was an order of magnitude smaller than divergence in lineages 2 to 4 (θ_w_ per site: 5.2e-5 to 7.2e-5, π per site: 4.5e-5 to 4.9e-5) and was highest in lineage 1 (θ_w_ per site: 2.3e-4; π per site: 2.1e-4) (Table 1). Tajima’s D was negative in all lineages, indicating an excess of low-frequency polymorphism, and values were closer to zero in lineages 1 and 4 (D=-0.56 and D=-0.82, respectively) than in lineages 2 and 3 (D=-1.45 and D=-1.72). The same differences in levels of variability across lineages, and individual summary statistics of the same order of magnitude, were observed if missing data were excluded from computations.

**Table 1.** Summary of population genomic variation in non-overlapping 100 kb windows Lineages 5 and 6 were not included in calculations because the sample sizes for these lineages were too small (*n*=2 and *n*=1, respectively); *n* is sample size; θ_w_ is Watterson’s θ per bp; π is nucleotide diversity per bp; H_e_ is haplotype diversity; K is the number of haplotypes; D is Tajima’s neutrality statistic.

| Lineage | $n$ | $S$ | $K$ | $H_e$ | $\theta_w$ | $\pi$ | $D$ |
| --- | --- | --- | --- | --- | --- | --- | --- |
| 1 | 10 | 57.6 | 3.6 | 0.31 | 2.25E-04 | 2.11E-04 | -0.558 |
| 2 | 14 | 19.7 | 7.2 | 0.20 | 6.94E-05 | 4.92E-05 | -1.454 |
| 3 | 16 | 21.5 | 7.9 | 0.17 | 7.24E-05 | 4.53E-05 | -1.718 |
| 4 | 6 | 10.5 | 4.3 | 0.38 | 5.15E-05 | 4.52E-05 | -0.824 |

Lineages 5 and 6 were not included in calculations because the sample sizes for these lineages were too small ( $n=2$ and $n=1$ , respectively); $n$ is sample size; $\theta_w$ is Watterson's $\theta$ per bp; $\pi$ is nucleotide diversity per bp; $H_e$ is haplotype diversity; $K$ is the number of haplotypes; $D$ is Tajima's neutrality statistic.

### Footprints of natural selection and the cost of pestification

We tested for standard neutral molecular evolution by the McDonald-Kreitman method, based on genome-wide patterns of synonymous and nonsynonymous variation (Table 2). The null hypothesis could be rejected for all four lineages (p<0.0001). The neutrality index, which quantifies the direction and degree of departure from neutrality, was greater than 1, indicating an excess of amino-acid polymorphism. This pattern suggests that lineages 1 to 4 accumulated slightly deleterious mutations during divergence from the Setaria-infecting lineage. Under near-neutrality, the ratio of nonsynonymous to synonymous nucleotide diversity π_N_π_S_ provides an estimate of the proportion of effectively neutral mutations that are strongly dependent on effective population size N_e_ [37]. The π_N_/π_S_ ratio ranged from 0.43 in lineage 1 to 0.61 in lineage 4, and was intermediate in lineages 2 and 3 (π_N_/π_S_=0.49), and the ratio of nonsense (i.e premature stop codons) to sense nonsynonymous mutations (P_nonsense_/P_sense_) followed the same pattern. Overall, the π_N_/π_S_ and P_nonsense_/P_sense_ ratios obtained suggest a higher proportion of slightly deleterious mutations segregating in lineage 4, and, to a lesser extent, in lineages 2 and 3, than in lineage 1. Assuming identical mutation rates, we can estimate that the long-term population size of lineage 1 (π_S_=0.00018/bp) was 2.5 to 3 times greater than that of the other lineages, consistent with the effect of N_e_ on the efficacy of negative selection predicted under near-neutrality.

**Table 2.**
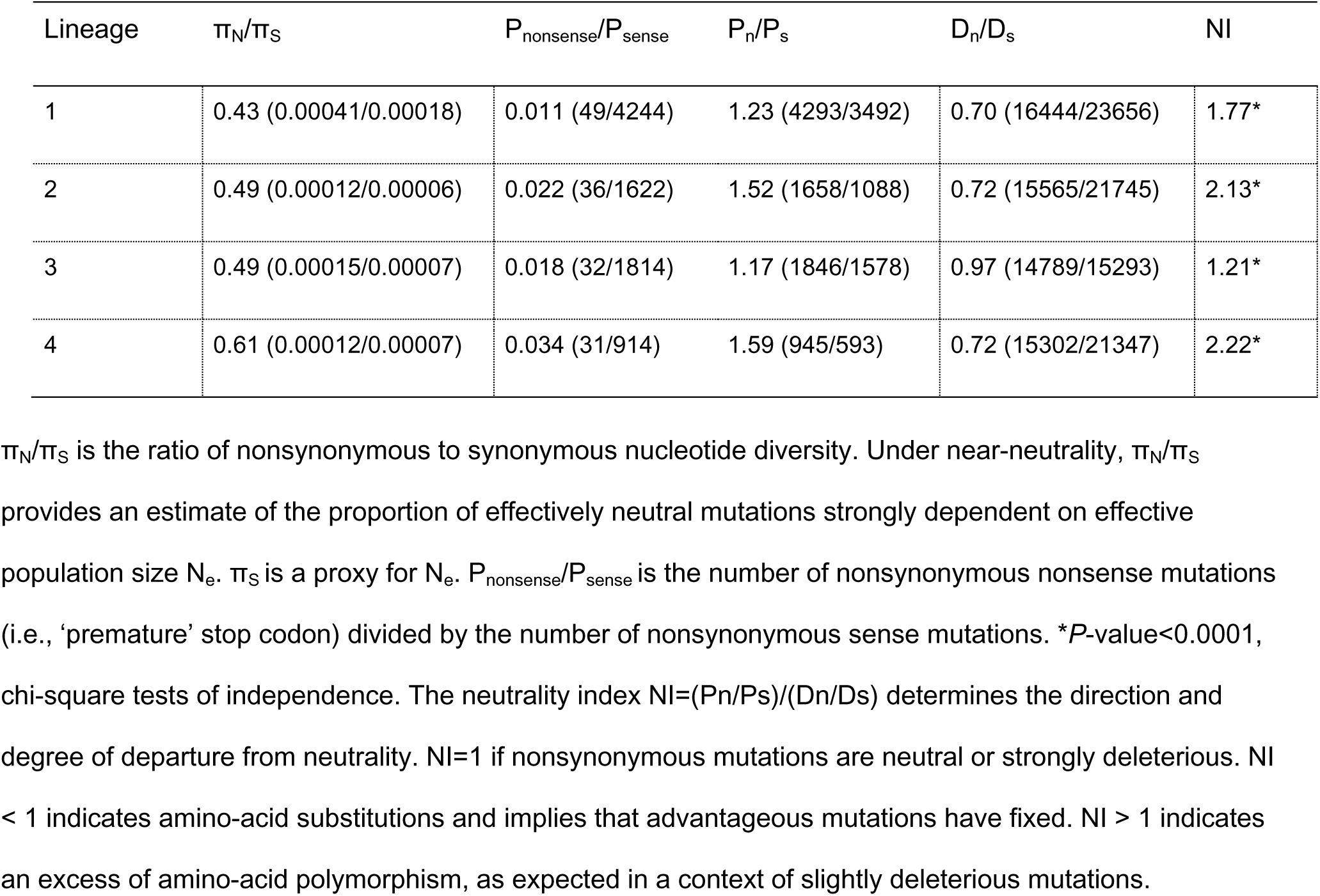
McDonald-Kreitman tests based on genome-wide patterns of synonymous and non synonymous variation, and measurements of the genome-wide intensity of purifying selection. Divergence was measured against predicted gene sequences of the Setaria-infecting *Magnaporthe oryzae* isolate US71.

### Distribution and reproductive biology of *M. oryzae* lineages

The strains of lineages 1 and 2 originated from rainfed upland rice, including rice grown in experimental fields. Lineage 2 was exclusively associated with tropical and temperate japonica, whereas lineage 1 was sampled from barley, tropical japonica and hybrid rice varieties (Table S1; Figure 1). Lineage 1 was restricted to continental Southeast Asia (Laos, Thailand, Yunnan). The reference laboratory strain GY-11 (=Guy11) was collected in French Guiana, from fields cultivated by Hmong refugees, who fled Laos in the 1970s. Lineage 2 was pandemic, and included all the European samples.

Lineage 3 and 4 samples originated from irrigated or rainfed upland/lowland rice. They were mostly associated with indica rice, with two samples collected from hybrid varieties and one collected from tropical japonica (Table S1; Figure 1). Lineage 3 was pandemic and included all sub-Saharan Africa samples, whereas lineage 4 was found on the Indian subcontinent, in Zhejiang (China) and the USA. Lineages 5 and 6 were collected from indica and tropical japonica varieties of rainfed upland rice in Yunnan and Hunan (China), respectively.

Lineages 2, 3 and 4 displayed low rates of female fertility (20%, 0%, 0%, respectively), and a significant imbalance in mating-type ratio (frequency of Mat-1: 100%, 14.3% and 100%, respectively; Chi^2^ test, *P*<0.001), whereas lineage 1 had a female fertility of 88.9% and a non-significant imbalance in mating type ratio (frequency of Mat-1:33.3%; Chi^2^ test, *P*=0.083). Lineage 5 was Mat-1 and only one of the two strains was female-fertile (no data for lineage 6).

### Pathogen compatibility range

Gallet et al. [38] analyzed the range of compatibility, in terms of the qualitative success of infection, between 31 *M. oryzae* isolates and 57 rice genotypes. Analyses of variance revealed a pattern of host-pathogen compatibility strongly structured by the host of origin of the isolates (i.e. the rice subspecies from which samples were collected). We investigated whether the compatibility between rice hosts and *M. oryzae* isolates was also structured by the lineage of origin of the isolates, by supplementing the dataset published by Gallet et al. [38] with pathotyping data for 27 isolates. We added microsatellite data to the SNP data, to overcome the absence of sequence data for 28 isolates, and we used clustering methods to confidently assign 46 of the 58 isolates with pathotyping data to identified lineages (no isolates could be assigned to lineage 5 or 6, see *Methods*). The final pathogenicity dataset included 46 isolates from lineages 1 to 4, inoculated onto 38 tropical japonica, temperate japonica, and indica varieties and 19 differential varieties with known resistance genes (Table S3).

Infection success (binary response) was analyzed with a generalized linear model. An analysis of the proportion of compatible interactions revealed significant effects of rice subspecies, pathogen lineage and the interaction between them (Table S4). The lineage effect could be explained by lineage 2 having a lower infection frequency than lineage 1 (comparison of lineages 1 and 2; z-value=-2.779; *p*-value=0.005) and lineage 3 having a higher infection frequency than lineage 1 (comparison of lineages 3 and 1; z-value=2.683; p-value=0.007), whereas the infection frequency of lineage 4 was not significantly different from that of lineage 1 (comparison of lineages 4 and 1; z-value=1.121; *p*-value=0.262). The rice subspecies effect could be attributed to tropical japonica varieties having a wider compatibility range than indica varieties (comparison of tropical japonica and indica; z-value=1.793; *p*-value=0.073), and temperate japonica having a wider compatibility range than indica varieties (comparison of temperate japonica and indica; *z*-value=1.830; *p*-value=0.067). The significant interaction between rice subspecies and pathogen lineage indicates that the effect of the lineage of origin of the isolate on the proportion of compatible interactions differed between the three rice subspecies. This interaction effect can be attributed to pathogen specialization on indica and tropical japonica, with lineage 1 (mostly originating from tropical japonica or from areas in which tropical japonica is grown) infecting tropical japonica varieties more frequently than indica varieties, lineage 2 (the lineage sampled from temperate japonica) infecting temperate japonica varieties more frequently than other varieties, lineages 3 and 4 (mostly originating from indica varieties) infecting indica varieties more frequently than tropical japonica varieties, and all four lineages infecting temperate japonica varieties at relatively high frequencies (Table S4; Figure 2A).

**Figure 2.**
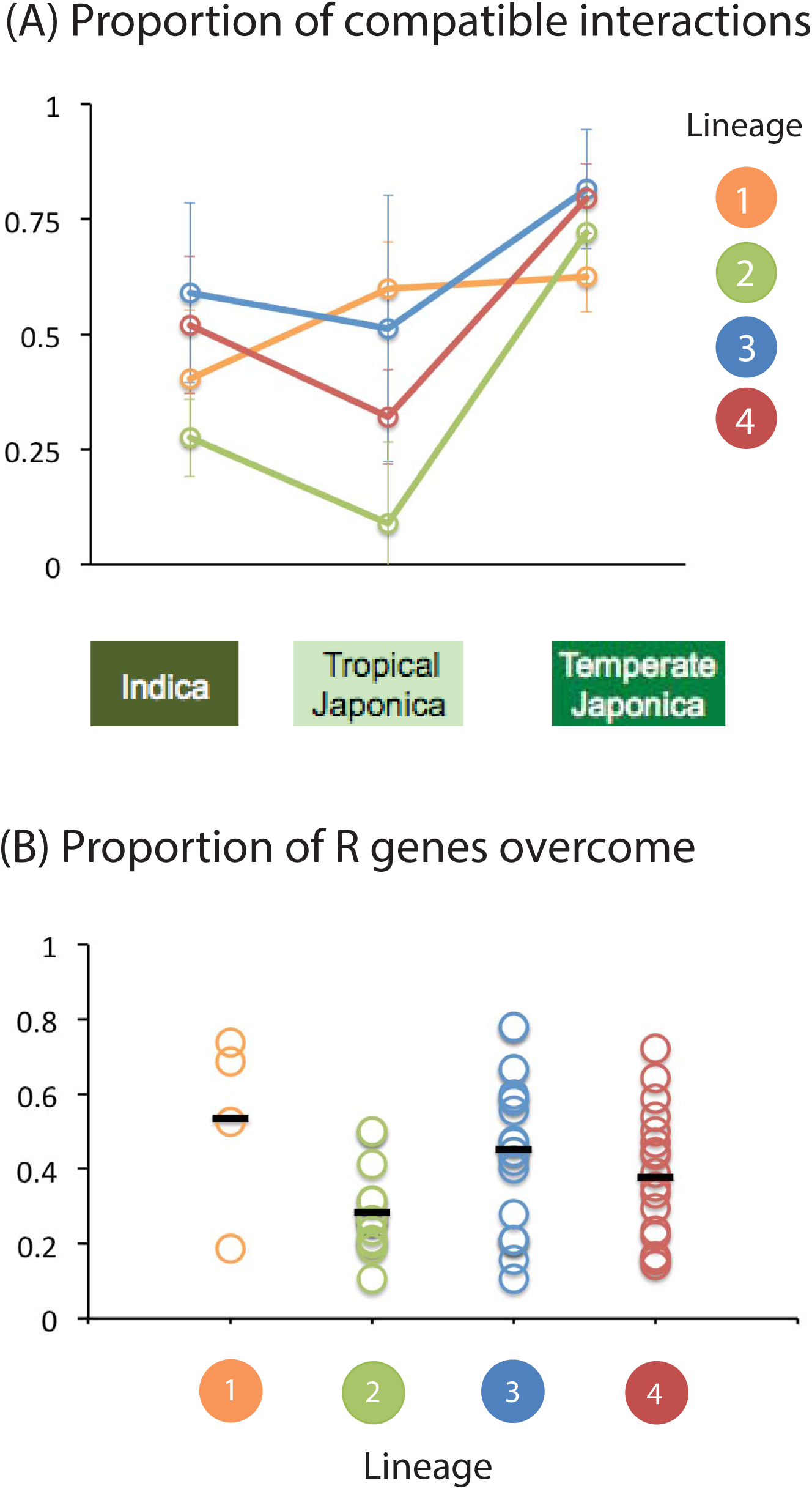
Proportion of compatible interactions between 46 isolates from lineages 1 to 4 of *M. oryzae* and 38 varieties representing three rice subspecies (B) and the proportion of R genes overcome by 36 isolates from lineages 1 to 4 of *M. oryzae* used to inoculate 19 differential lines of rice.

Major resistance (R) genes can be a major determinant of pathogen host range, and promote divergence between pathogen lineages by exerting strong divergent selection on a limited number of pathogenicity-related genes [39-41]. We investigated the possible role of major resistance (R) genes in the observed differences in compatibility between rice subspecies and pathogen lineages, by challenging 19 differential varieties with the 46 isolates assigned to lineages 1 to 4. An analysis of the number of R genes overcome revealed a significant effect of pathogen lineage (Table S5). This effect was driven mostly by lineage 2, which overcame fewer R genes than the other lineages (Table S5; Figure 2B).

### Recombination within and between lineages

We visualized evolutionary relationships, while taking into account the possibility of recombination within or between lineages, using the phylogenetic network approach Neighbor-Net, as implemented in Splitstree 4.13 [42]. Neighbor-Net is an agglomerative method that generates planate split graph representations. A *split* is a partitioning of the dataset, and a collection of splits is considered *compatible* if they fall within the set of splits of a tree. Gene genealogies represent compatible collections of splits, whereas Neighbor-Net can be used to visualize conflicting phylogenetic signals, represented by network reticulation, through a condition weaker than compatibility. The Neighbor-Net network inferred from the set of 16,370 SNPs without missing data presented a non tree-like structure of the inner connections between lineages, consistent with genetic exchanges between unrelated isolates or incomplete lineage sorting (Figure 3). Greater network reticulation was observed between lineages 1, 5 or 6 and the other lineages than between these other lineages themselves. Lineages 2 to 4 had long interior branches and star-like topologies, consistent with long-term clonality.

**Figure 3.**
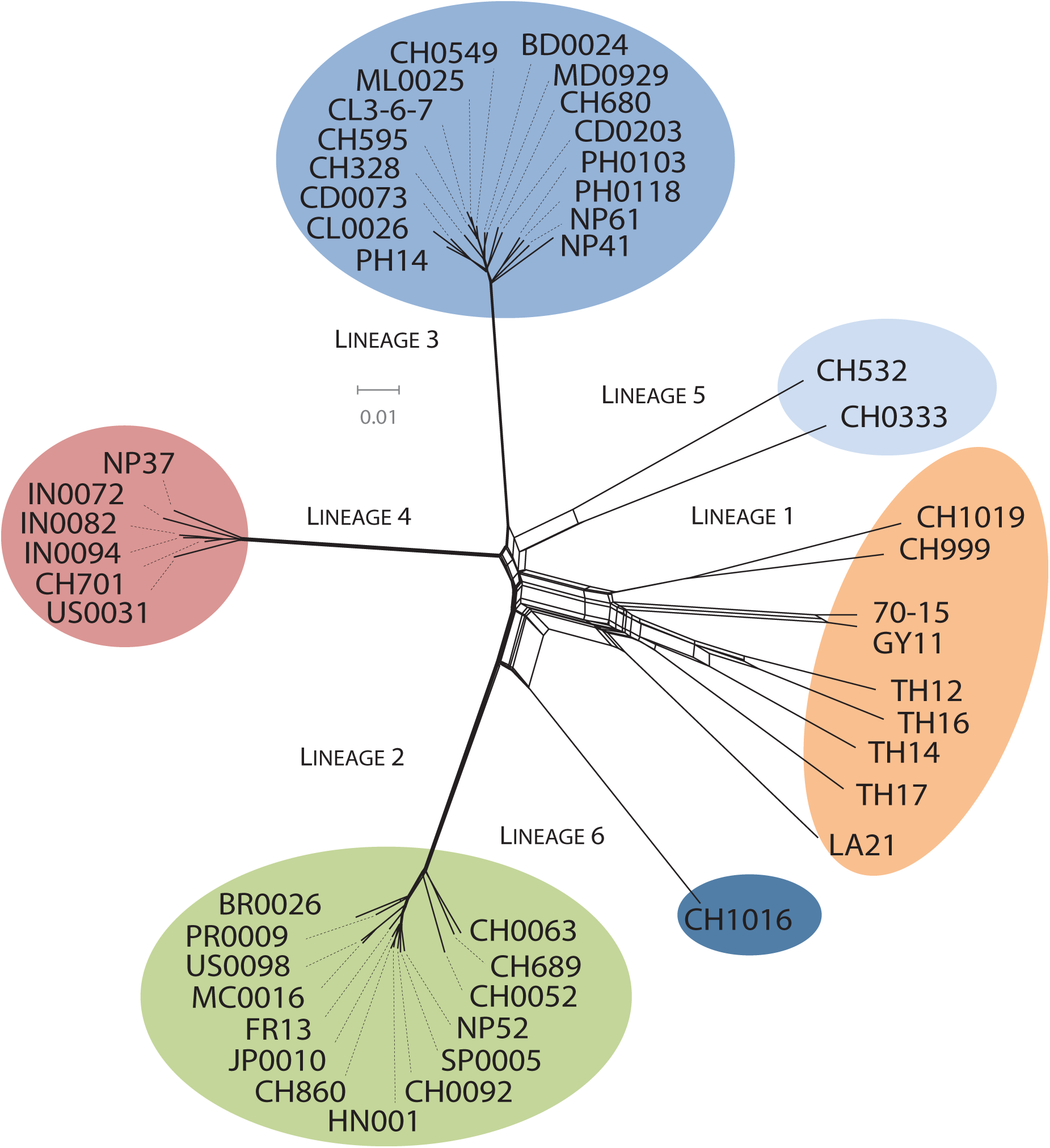
Neighbor-Net networks showing relationships between haplotypes identified on the basis of the full set of 16,370 SNPs without missing data, (A) in the whole sample set, (B) in lineage 1, (C) in lineage 2, (D) in lineage 3, (E) in lineage 4.

We evaluated the amount of recombination within lineages, by estimating the population recombination parameter (rho=2N_e_r) and testing for the presence of recombination with a likelihood permutation test implemented in the pairwise program in LDHat. Recombination analyses confirmed the heterogeneity between lineages of the contribution of recombination to genomic variation, with recombination rates averaged across chromosomes being more than two to three orders of magnitude higher in lineage 1 (10.57 crossovers/Mb/generation) than in other lineages (lineage 2: 0.28; lineage 3: 0.01; lineage 4: 0.33 crossovers/Mb/generation) (Table 3). Splitstree analyses displaying the reticulations within each lineage and testing for recombination with the PHI test were consistent with this pattern (Table 3; Figure 3; Figure S1). The null hypothesis of no recombination was rejected only for lineages 1 and 4 [43].

**Table 3.** Estimates of the population recombination rate ρ, tests of recombination based on homoplasy and linkage disequilibrium, proportion of homoplastic SNPs

| Lineage | ρ (crossovers/Mb/generation) |  |  |  |  |  |  |  | Homoplastic<br>SNPs | PHI test<br>(*p*-value) |
| --- | --- | --- | --- | --- | --- | --- | --- | --- | --- | --- |
|  | Chromosomes |  |  |  |  |  |  |  |  |  |
|  | 1 | 2 | 3 | 4 | 5 | 6 | 7 | Mean |  |  |
| 1 | 8.6* | 3.8* | 15.1* | 1.4 | 8.5* | 10.6* | 13.5* | 10.57 | 34.56 % | 0.0000 |
| 2 | 0.0 | 0.2 | 0.2* | 0.3 | 0.6 | 0.5 | 0.3 | 0.28 | 0.09 % | 0.0944 |
| 3 | 0.4* | 0.2* | 0.0 | 0.0 | 0.0 | 0.0 | 0.0 | 0.01 | 0.47 % | 0.0535 |
| 4 | 0.2 | 0.2* | 0.3 | 0.4 | 0.4 | 0.4 | 0.4 | 0.33 | 0.40 % | 0.0014 |

\* $P < 0.05$ . PHI test assesses pairwise homoplasy. The null hypothesis of no recombination was tested, for the PHI test and for $\rho$ , using random permutations of the positions of the SNPs based on the expectation that sites are exchangeable if there is no recombination. For the $\rho$ test, significance was determined from the distribution of maximum composite likelihood values calculated from permuted data.

Differences in recombination-based variation between lineages were confirmed by analyses of homoplasy (Table 3). Homoplastic sites display sequence similarities that are not inherited from a common ancestor, instead resulting from independent events in different branches. Homoplasy can result from recurrent mutations or recombination, and the contribution of recombination to homoplasy is expected to predominate in outbreeding populations. Homoplastic sites were identified by mapping mutations onto the total-evidence genome genealogy with the ‘Trace All Characters’ function of Mesquite [44], applying ancestral reconstruction under the maximum parsimony optimality criterion. The resulting matrix of ancestral states for all nodes was then processed with a python script, to determine the number of mutations that had occurred at each site within each lineage, counting sites displaying multiple substitutions across the tree as homoplastic. Only 0.09%, 0.47% and 0.40% of the SNPs were homoplastic in lineages 2, 3 and 4, respectively, versus 34.6% in lineage 1 (lineages 5 and 6 were not tested due to the small sample size). The very small numbers of homoplastic sites in lineages 2, 3 and 4 suggest that these lineages are largely clonal, whereas the high level of homoplasy detected in lineage 1 is consistent with repeated recombination events between the strains of this lineage.

We assessed the genomic impact of recombination, by analyzing patterns of linkage disequilibrium (LD), corresponding to the tendency of different alleles to occur together in a non-random manner. For lineage 1 (S=13 kSNPs), LD decayed smoothly with physical distance, reaching half its maximum value at about 10 kb, whereas, for lineages 2, 3 and 4 (S=3.7, 3.2 and 2.7 kSNPs), no LD decay pattern was observed (Figure S2). These analyses also revealed that background LD levels were no higher in lineages 2, 3, or 4, which appeared to be largely clonal, than in lineage 1. However, both simulation work and empirical data have shown that population history, including bottlenecks and admixture, strongly affects the background level of LD in the population [45].

### Genome scan for genetic exchanges between lineages

We scanned the genomes for the exchange of mutations between lineages, using a method based on lineage-diagnostic SNPs and a probabilistic method of ‘chromosome painting’ (Figure 4). In the lineage-diagnostic SNP approach, each isolate is removed from the dataset in turn to identify SNPs specific to a particular lineage (i.e. biallelic sites displaying a mutation specific to a given lineage). Each focal isolate is then added back to the dataset and scanned for the presence of lineage-diagnostic SNPs identified in lineages other than its lineage of origin. Using this approach, we identified 515 lineage-diagnostic singletons with 276, 96 and 140 singletons in lineages 1, 5 and 6, respectively, and only one singleton in lineages 2, 3 and 4. Putatively migrant singletons were assigned to all other lineages for lineages 1 and 5, and to all other lineages except lineage 1 for lineage 6 (Table S6). ‘Chromosome painting’ is a probabilistic method for reconstructing the chromosomes of each individual sample as a combination of all other homologous sequences. We identified the migrant mutations present in each isolate, these mutations being defined as having a probability greater than 90% of having been copied from a lineage other than the lineage of origin of the focal isolate. This method uses population data from recipient populations only, and we were therefore able to include only lineages 1 to 4 in the analysis. Chromosome painting identified 464 migrant mutations, all of which segregated in lineage 1. Putative migrant mutations were assigned to all five of the other lineages (92.8 mutations per lineage, on average), with lineage 2 making the largest contribution (165 mutations) and lineage 4 the smallest contribution (39 mutations).

**Figure 4.**
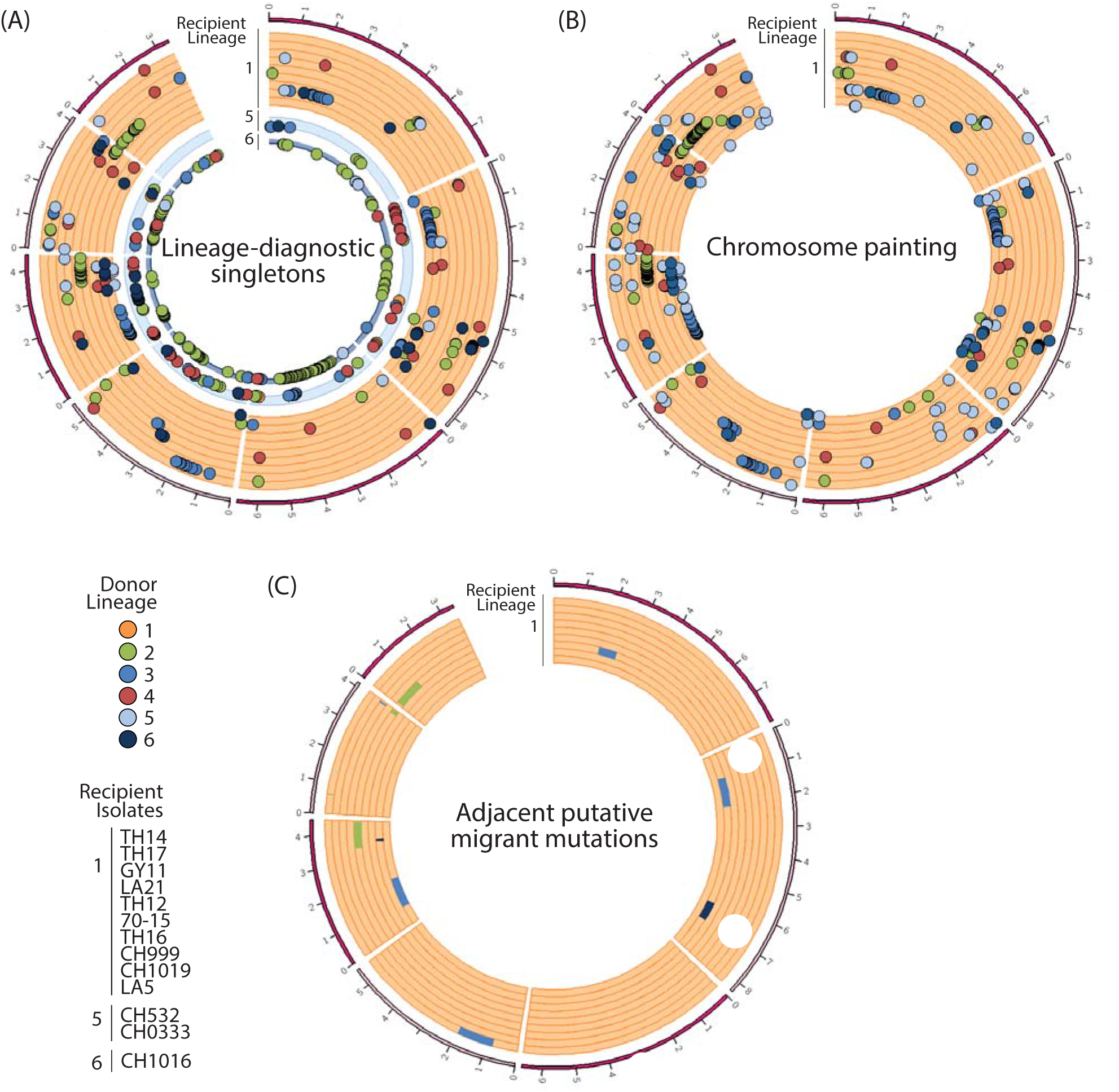
Genomic distribution of candidate immigrant mutations in lineages 1, 5 and 6. (A) Lineage-diagnostic mutations segregating as singletons in other lineages. (B) Lineage 1 mutations for which the most probable donor is lineage 2, 3, or 4 in probabilistic chromosome painting analysis. Lineages 5 and 6 (*n*=2 and *n*=1, respectively) could not be included as recipient populations in the chromosome painting analysis due to their small sample sizes. No candidate immigrant mutations were identified in lineages 2, 3, and 4. (C) Genomic regions corresponding to series of adjacent putative migrant mutations identified with lineage-diagnostic singletons in lineage 1. Chromosomes 1 to 7, in clockwise order, with ticks at megabase intervals.

The sets of putative migrant mutations identified by the two methods matched different sets of genes enriched in NOD-like receptor [46], HET-domain [47] or GO term ‘lipid catabolic process’ (Table S6) genes. However, the presence of false positives due to the random sorting of ancestral polymorphisms in lineage 1 and other lineages cannot be excluded. We minimized the impact of the retention of ancestral mutations, by reasoning that series of adjacent mutations are more likely to represent genuine gene exchange events. We identified all the genomic regions defined by three adjacent putative migrant mutations originating from the same donor lineage. We searched for such mutations among the set of putative migrant mutations identified by the two methods. We identified 12 such regions in total, corresponding to 1917 genes. Functional enrichment tests for each recipient isolate revealed enrichment in the GO term ‘pathogenesis’ for isolate CH999, the GO term ‘phosphatidylinositol biosynthetic process’ for isolate TH17 and the GO term ‘telomere maintenance’ for isolate CH1019 (Table S6).

### Molecular dating

We investigated the timing of rice blast emergence and diversification, by performing Bayesian phylogenetic analyses with Beast. Isolates were collected from 1967 to 2009 (Table S1), making it possible to use a tip-based calibration approach to estimate evolutionary rates and ancestral divergence times together. We analyzed the linear regression of sample age against root-to-tip distance (i.e. the number of substitutions separating each sample from the hypothetical ancestor at the root of the tree). The temporal signal obtained in this analysis was strong enough for thorough tip-dating inferences (Figure S3) [48]. We therefore used tip-dating to estimate the rate at which mutations accumulate (i.e. the substitution rate) and the age of every node in the tree, including the root (i.e. time to the most recent common ancestor), simultaneously. At the scale of the genome, the mean substitution rate was estimated at 1.98e-8 substitutions/site/year (Figure S3). The six rice-infecting lineages were estimated to have diversified ~900 to ~1300 years ago (95% HPD [175-2700] years ago) (Figure 5). Bootstrap node support was strong and similar node age estimates were obtained when the recombining lineage 1 and the potentially recombining lineages 5 and 6 (not shown) were excluded, indicating the limited effect of recombination on our inferences. We also inferred that the ancestor of rice-infecting and Setaria-infecting lineages lived ~9,800 years ago. However, the credibility intervals were relatively large (95% HPD [1200-22,000] years ago), covering the period from japonica rice domestication and Setaria domestication to the last glacial maximum, and overlapping with previous estimates suggesting that the rice‐ and Setaria-infecting lineages diverged shortly after rice domestication, or even during the period of rice domestication (range of point estimates in ref [15]: 2500 to 7300 years ago).

**Figure 5.**
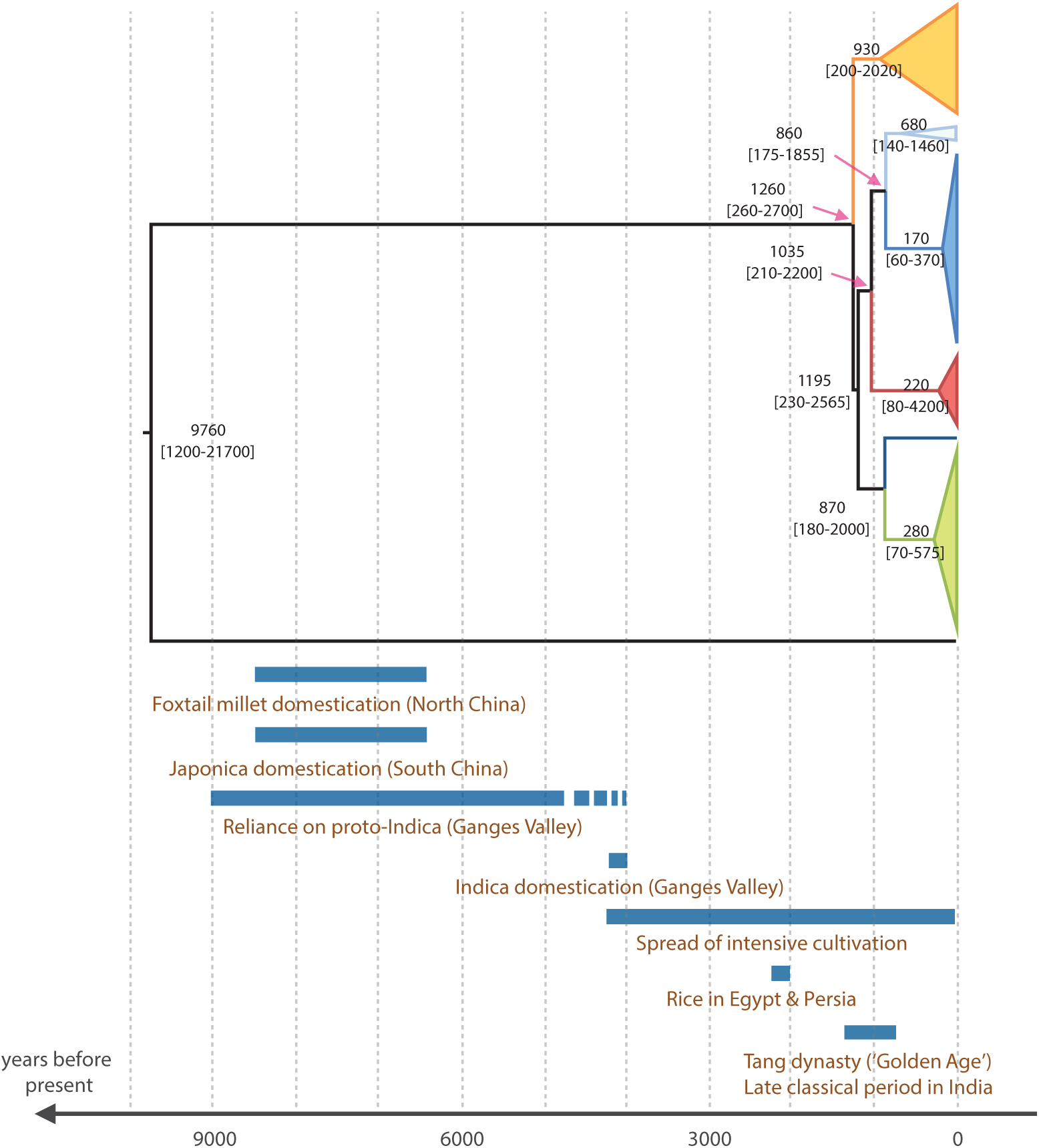
Tip-calibrated genealogy inferred by maximum-likelihood phylogenetic inference in Beast 1.8.2, based on single-nucleotide variation in 50 *M. oryzae* genomes. Approximate historical periods are shown for context.

## Discussion

We performed a whole-genome sequence analysis of 50 isolates with different temporal and spatial distributions, to elucidate the emergence, diversification and spread of *M. oryzae* as a rapidly evolving pathogen with a devastating impact on rice agrosystems. Analyses of population subdivision confirmed the four lineages previously identified by Saleh et al. [21]. Previous analyses of microsatellite data were unable to resolve the genealogical relationships between clusters or to capture the phylogenetic depth of population subdivision within *M. oryzae.* By contrast, our population genomic analyses of resequencing data revealed weak divergence between clusters (absolute divergence dxy of the order of 10^-4^ differences per base pair), consistent with recent diversification. Phylogenetic analyses using sampling dates for calibration confirmed the recent origin of the six lineages, with estimates of divergence time ranging from ~900 to ~1300 years ago (95% credible intervals [175-2700] years ago). Lineage 1 (which includes the reference strains GY11 and 70-15) was found in mainland Southeast Asia and originates from barley, tropical japonica or undetermined varieties. All isolates from lineages 1, 5 and 6 were collected in rainfed upland agrosystems typical of japonica rice cultivation, and pathogenicity test results were consistent with the local adaptation of lineage 1 to tropical japonica rice. Lineage 2 was pandemic in irrigated fields of temperate japonica rice outside Asia, and cross-inoculation experiments revealed specialization on this host and an ability to overcome fewer R genes, on average, than other lineages. Lineages 3 and 4 were associated with indica. Lineage 3 is pandemic, and crossinoculation indicated local adaptation to this host, relative to tropical japonica, although lineages 3 and 4 had relatively wide compatibility ranges consistent with generalism. One possible explanation of the wide compatibility range of temperate japonica varieties and the narrow compatibility range of lineage 2 is that temperate japonica varieties have smaller repertoires of R genes, as resistance to blast is of less concern to breeders in temperate irrigated conditions, which are less conducive to epidemics [38].

The continental Southeast Asian lineage was the most basal in total-evidence genome genealogies, reflecting a pathway of domesticated Asian rice evolution [16, 18] in which the *de novo* domestication of rice occurred only once, in japonica. However, the diversification of *M. oryzae* into multiple rice-infecting lineages (point estimates ranging from ~900 to ~1300 years ago) appears to be much more recent than the *de novo* domestication of rice (8500-6500 years ago [16,49,50]), the spread of rice cultivation in paddy fields, and the domestication of indica in South Asia, following introgressive hybridization from the early japonica gene pool into ‘proto-indica’ rice (about 4000 years ago [16, 51]). At the time corresponding to the upper bound of the 95% credible interval (2700 years ago), japonica rice and paddy-field cultivation had spread to most areas of continental and insular South, East and Southeast Asia, and indica rice was beginning to spread out of the Ganges plains [16, 52]. The point estimates for the splitting of *M. oryzae* lineages correspond to the Tang dynasty (‘the Golden Age’) in China, and the late classical period in India, during which food production became more rational and scientific and intensive irrigated systems of cultivation were developed, bringing about economic, demographic and material growth [53].

Genome scans based on polymorphism and divergence revealed heterogeneity in the genomic and life-history changes associated with the emergence and spread of the different lineages. Using microsatellite data and a larger collection of samples, Saleh et al. [21] identified differences in variability between lineages, with similar or higher levels of genetic variability in lineages 1 and 4 than in lineages 2 and 3. Lineages 1 and 4 were also the only lineages displaying biological features (fertile female rates and mating type ratios) consistent with sexual reproduction. Our genome-wide analyses of variability and linkage disequilibrium provided clear evidence that the continental Southeast Asian lineage 1 displays recombination and is genetically diverse, suggesting that sexual reproduction occurs and that long-term population size is relatively high, whereas pandemic lineages 2 and 3 are largely clonal and genetically depauperate, suggesting a lack of sexual reproduction and demographic bottlenecks associated with their emergence in agrosystems. However, population genomic analyses did not confirm the previously reported high variability and capacity for sexual recombination of the South Asia/US lineage 4 [21], possibly due to differences in sample size between studies. The null hypothesis of clonality was not rejected in PHI-tests for recombination, but both total (θ_w_) and average (π) nucleotide diversity, and the population recombination rate (ρ), were of the same order of magnitude in lineage 4 as in lineages 2 and 3, consistent with a lack of recombination and a small effective population size.

The patterns of polymorphism and diversity at non-synonymous and synonymous sites indicated that deleterious mutations were particularly abundant in clonal lineages 2-4 of *M. oryzae*, with their smaller long-term population size, consistent with a higher cost of pestification in these lineages. The introgression of genetic elements from clonal lineages harboring greater loads of deleterious mutations may counteract the efficient purging of deleterious mutations in the recombining lineage 1 from mainland Southeast Asia, and lead to smaller differences in the proportion of nonsynonymous mutations between recombining and clonal lineages. However, the extensive variability of the origin and genomic distribution of the detected putative migrant mutations suggests that most of these mutations are false-positive, with only series of adjacent mutations of this type originating from the same donor lineage corresponding to genuine genetic exchange events. Field-scale studies in areas in which different lineages coexist should provide more detailed insight into the relative importance of interlineage recombination, and make it possible to determine whether genetic exchanges are driven by positive selection or are an incidental byproduct of the sympatric coexistence of interfertile lineages. We hypothesize that the accumulation of deleterious mutations in pandemic clonal complexes and gene flow into sexual lineages during disease emergence and spread are widespread phenomena, not due to idiosyncrasies of *M. oryzae*, and we expect these patterns to hold true in other invasive fungal plant pathogens.

An examination of additional isolates from under-sampled geographic regions (including Africa and South America), based on sequencing approaches and sampling schemes tailored to detect adaptation from *de novo* mutations, will be required to enhance our understanding of the biogeography of *M. oryzae* and the genetic basis of adaptation in the different *M. oryzae* lineages. Nevertheless, the catalog of variants detected in our study provides a solid foundation for future research into the population genomics of adaptation in *M. oryzae.* Our work also provides a population-level genomic framework for defining molecular markers for the control of rice blast and investigations of the molecular basis of the differences in phenotype and fitness between divergent lineages.

## Methods

### Genome sequencing and SNP calling

Sequencing libraries were prepared and Illumina HiSeq 2500 sequencing was performed either at Beckman Coulter Genomics (Danvers, USA) or at the Iwate Biotechnological Research Center (Table S1). Genomic DNA for sequencing at BCG was isolated from 100 mg of fresh mycelium grown in liquid medium. The mycelium was treated with enzymes degrading the cell walls (mainly betaglucanase) and then incubated in lysis buffer (Triton 2× – 1% SDS - 100 mM NaCl – 10 mM Tris-HCl – 1 mM EDTA). Nucleic acids were extracted by treatment with chloroform:isoamyl alcohol (24:1), followed by precipitation overnight in isopropanol. They were then rinsed in 70% ethanol. The nucleic acid extract was treated with RNase A (0.2 mg/mL final concentration) to remove RNA. The DNA was purified by another round of chloroform:isoamyl alcohol (24:1) treatment. Genomic DNA for sequencing at IBRC was isolated with a protocol adapted from the animal tissue (Mouse tail) protocol available in the Promega Wizard^®^ Genomic DNA Purification Kit. Nucleic acids were extracted from 20 mg of fresh mycelium grown in liquid medium, which was ground into powder in liquid nitrogen, with a pre-chilled pestle and mortar. The centrifugation time specified in the mouse tail protocol was increased to 15 min, and centrifugation was carried out at 4°C, after precipitation for 3 hours at - 20°C. Nucleic acids were resuspended in water, treated with RNase A (0.2 mg/mL final concentration), purified by treatment with chloroform:isoamyl alcohol (24:1), precipitated overnight in isopropanol supplemented with 0.1 volumes of sodium acetate (3 M pH = 5), and rinsed in 70% ethanol.

Sequencing reads were either paired-end (read length 100 nucleotides, insert size ~ 500 bp, DNAs sequenced at IBI) or single-end (read length 100 nucleotides, DNAs sequenced by BCG). Reads were trimmed to remove barcodes and adapters, and were then filtered to eliminate sequences containing ambiguous base calls. Reads were mapped against the 70-15 reference genome version 8 [10] with bwa [54] (subcommand alb, option ‐n 5; subcommand sampe option-a 500). Alignments were sorted with SAMTOOLS [55], and reads with a mapping quality below 30 were removed. Duplicates were removed with PICARD (http://broadinstitute.github.io/picard/). We used realigner-targetcreator, targetcreator and indelrealigner within the genome analyses toolkit (GATK) [56] to define intervals to target for local realignment and for the local realignment of reads around indels, respectively, and unified genotyper to call SNPs. We used GATK’s SelectVariants to apply hard filters and to select high-confidence SNPs based on annotation values. Numbers of reference and alternative alleles were calculated with JEXL expressions based on the vc.getGenotype().getAD() command. Variants were selected based on the following parameters: counts of all reads with a MAPQ = 0 below 3.0 (MQ0 in GATK), number of reference alleles + number of alternative alleles ≥15.0, and number of reference alleles/number of alternative alleles ≤ 0.1. With these parameters, SNP calls are limited to positions with relatively high sequencing depths and limited discordance across high-quality sequencing reads. We used a second SNP caller, Freebayes v0.9.10-3-g47a713e [57], to assess the impact of the SNP calling method on the sets of SNPs detected, given the presence in our dataset of isolates sequenced at relatively low depth (<10X). We set the ‐‐min-alternate-count option to one in Freebayes. When the sample-by-sample Freebayes SNP calls were compared with the GATK SNP calls, after filtration, Freebayes identified 1.63X (stdev 0.28) more SNPs per sample on average than analyses with GATK, and 92.3% (stdev 2.3) of the SNPs identified with GATK were also identified with Freebayes. The size of the intersection between the sets of SNPs identified by the two methods was negatively correlated with sequencing depth (i.e. the concordance between SNP callers was higher for isolates sequenced less deeply), indicating a minimal impact of isolates sequenced at lower depth on confidence in SNP calls. When the multisample Freebayes SNP calls were compared with the GATK SNP calls, after filtration, 83% of the SNPs identified with GATK were confirmed with Freebayes, and the GATK SNPs that were not confirmed with Freebayes were identified in sets of isolates with a genome-wide sequencing depth of 47.8X on average (stdev 8.2), consistent with a minimal impact of isolates sequenced at lower depth on confidence in SNP calls. High-confidence SNPs were annotated with SnpEff v4.3 [58].

### Mating type and female fertility assays

Mating type and female fertility had previously been determined [23], or were determined as previously described [59].

### Genealogical relationships and population subdivision

Total-evidence genealogy was inferred with RAxML from pseudo-assembled genomic sequences (i.e. tables of SNPs converted into a fasta file using the reference sequence as a template), assuming a general time-reversible model of nucleotide substitution with the Γ model of rate heterogeneity. Bootstrap confidence levels were determined with 100 replicates. DAPC was performed with the Adegenet package in R [60]. Sites with missing data were excluded. We retained the first 20 principal components, and the first six discriminant functions.

### Diversity and divergence

Polymorphism and divergence statistics were calculated with Egglib 3.0.0b10 [61], excluding sites with >30% missing data. The neutrality index was calculated as (P_n_/P_s_)/(D_n_/D_s_), where P_n_ and P_s_ are the numbers of nonsynonymous and synonymous polymorphisms, and D_n_ and D_s_ are the numbers of nonsynonymous and synonymous substitutions, respectively. D_n_ and D_s_ were calculated with Gestimator [62] using the Setaria-infecting lineage as an outgroup. P_n_ and P_s_ were calculated with Egglib.

### Linkage disequilibrium and recombination

The coefficient of linkage disequilibrium (r^2^) [63] was calculated with Vcftools [64], excluding missing data and sites with minor allele frequencies below 10%. For all lineages, we calculated r^2^ between all pairs of SNPs less than 100 kb apart and averaged LD values in distance classes of 1 kb for lineages 1 and 4, and 10 kb for lineages 2 and 3,to minimize noise due to low genetic diversity. Only sites without missing data and with a minor allele frequency above 10% were included, to minimize the dependence of r^2^ on minor allele frequency [65]. Recombination rates were estimated for each chromosome with pairwise in LDhat version 2.2 [66]. Singletons and sites with missing data were excluded.

### Pathogenicity tests

We used pathotyping data for 31 isolates previously described by Gallet et al. [38]. We supplemented this dataset with pathotyping data for 27 isolates, produced by the same authors, using the same protocol, but not included in the publication due to uncertainty in the nature of the rice subspecies of origin. We used a combination of multilocus microsatellite and SNP data to assign the 58 pathotyped isolates to the six lineages, because SNP data were available for only 30 pathotyped isolates (20 of the 31 isolates from Gallet et al. and 10 of the 27 additional isolates). Multilocus microsatellite genotypes at 12 loci were obtained from the Saleh et al. [21] dataset, or produced as described by Saleh et al. [21]. We improved the accuracy of assignment tests by adding the 19 isolates that had been sequenced but for which no pathotyping data were available to the full dataset, which included 77 multilocus genotypes in total (58 pathotyped isolates, and 19 additional non-pathotyped isolates). For 49 of the 77 isolates for which genomic data were available, we retained 1% of the SNP loci with no missing data (i.e. 164 SNPs). Missing data were introduced at SNP and microsatellite loci for the 28 non-sequenced isolates and the four sequenced isolates without microsatellite data.

structure 2.3.1 program was used for assignment [67-69]. The model implemented allowed admixture and correlation in allele frequencies. Burn-in length was set at 10 000 iterations, and the burn-in period was followed by 40,000 iterations. Four independent runs were performed to check for convergence. At *K*=6 the four main clusters identified with the full genomic dataset were recovered, although 15 of the 77 genotypes could not be assigned due to admixture or a lack of power. Finally, 46 of the 58 isolates inoculated could be assigned to lineages 1 to 4; the other 12 isolates could not be assigned to a specific lineage among lineages 1, 5 and 6, and were not analyzed further (Figure S4). Infection success was analyzed with a generalized linear model with a binomial error structure and logit link function. Treatment contrasts were used to assess the specific degrees of freedom of main effects and interactions.

### Genome scan for genetic exchanges

Probabilistic chromosome painting was carried out with Chromopainter version 0.0.4 [70]. This method ‘‘paints’’ individuals in ‘‘recipient’’ populations as a combination of segments from ‘‘donor’’ populations, using linkage information for probability computation, assuming that linked alleles are more likely to be exchanged together during recombination events. All lineages were used as donors, but only lineages 1-4 were used as recipients (sample size too small for lineages 5 and 6). We initially ran the model using increments of 50 expectation-maximization iterations, starting at 10 iterations, and examined the convergence of parameter estimates to determine how many iterations to use. Hence, the recombination scaling constant N_e_ and emission probabilities (μ) were estimated in lineages 1-4 by running the expectation-maximization algorithm with 200 iterations for each lineage and chromosome. Estimates of N_e_ and μ were then calculated as averages weighted by chromosome length (N_e_=8160 for all lineages, lineage 1; μ=0.0000506; lineage 2, μ=0.0000171; lineage 3, μ=0.000021; lineage 4, μ=0.000011). These parameter values and the per-chromosome recombination rates estimated with LDHAT were then used to paint the chromosome of each lineage, considering the remaining lineages as donors, using 200 expectation-maximization iterations. We used a probability threshold of 0.9 to assign mutations in a recipient lineage to a donor lineage.

### Tip-calibrated phylogenetic analysis

Tip-calibrated phylogenetic inferences were performed with only the 48 isolates for which sampling date was recorded, i.e. all isolates except the reference 70-15 and PH0018 isolates, with the exclusion of missing data. We investigated whether the signal obtained with our dataset was sufficiently high for thorough tip-dating inferences, by building a phylogenetic tree with PhyML [71], without constraining tip-heights on the basis of isolate sampling time, and then fitting root-to-tip distances (a proxy for the number of substitutions accumulated since the most recent common ancestor, TMRCA) to collection dates with TempEst [70]. We observed a significant positive correlation (Figure S3), demonstrating that the temporal signal was sufficiently strong for thorough tip-dating inferences at this evolutionary scale. The tip-calibrated inferences were then carried out using Markov chain-Monte Carlo sampling in Beast 1.8.2 [72]. The topology was fixed as the total-evidence genome genealogy inferred with RAxML. We used an annotation of the SNPs with SNPEff [57] to partition Bayesian inference (i.e. several substitution models and rates of evolution were fitted to the different sets of SNPs during a single analysis). The optimal partitioning scheme and the best-fit nucleotide substitution model for each partitioning of the genome were estimated with PartitionFinder software [73]. The best partitioning was obtained for *K*=3 schemes (synonymous: HKY, non-synonymous: GTR and non-exonic SNPs: GTR) and was used for subsequent analyses. Node age was then estimated with this optimal partitioning scheme. Rate variation between sites was modeled with a discrete gamma distribution, with four rate categories. We assumed an uncorrelated lognormal relaxed clock, to account for rate variation between lineages. We minimized prior assumptions about demographic history, by adopting an extended Bayesian skyline plot approach, to integrate data over different coalescent histories. The tree was calibrated using tip-dates only. We applied flat priors (i.e., uniform distributions) for substitution rate (1 x 10^-12^ – 1 x 10^-2^ substitutions/site/year) and for the age of any internal node in the tree (including the root). We ran five independent chains, in which samples were drawn every 5,000 MCMC steps, from a total of 50,000,000 steps, after a discarded burn-in of 5,000,000 steps. We checked for convergence to the stationary distribution and for sufficient sampling and mixing, by inspecting posterior samples (effective sample size >200). Parameter estimation was based on samples combined from the different chains. The best-supported tree was estimated from the combined samples using the maximum clade credibility method implemented in TreeAnnotator.

### Functional enrichment

Gene enrichment analysis was conducted with the R package topGO for GO terms, and Fisher’s exact tests for enrichment in HET-domain genes, NLRs, small-secreted protein and MAX-effector genes. MAX-effector genes were obtained from de Guillen et al. [71], NLRs were as identified in Dyrka et al. [46], and small secreted proteins and HET-domain proteins were identified with Ensembl’s Biomart.

## Acknowledgments

We thank the scholars who contributed samples, François Bonnot and Romain Gallet for assistance with statistics, and the Southgreen and Migale computing facilities.

## Supporting information

**Figure S1.** Neighbor-Net networks showing relationships between haplotypes identified on the basis of the full set of 16,370 SNPs without missing data, (A) in lineage 1, (B) in lineage 2, (C) in lineage 3, (D) in lineage 4.

**Figure S2.** Linkage disequilibrium (r^2^) against distance. Only SNPs of less than 100 kb and with a minor allele frequency of more than 10% are shown. Averaged values in 1 kb windows for lineages 1 and 4. Averaged values in 10 kb windows for lineages 2 and 3, to minimize the noise associated with smaller sample size.

**Figure S3.** (A) Root-to-tip distances (mutations/site) estimated with Beast are correlated with collection date; (B) Marginal posterior densities of the substitution rates of the three data partitions (non-coding sites, nonsynonymous sites and synonymous sites), as estimated by tip-calibrated phylogenetic analysis.

**Figure S4.** Proportions of ancestry in *K*=6 ancestral populations inferred with STRUCTURE program from 77 multilocus genotypes. Individuals were genotyped at 12 microsatellite loci and 164 SNP loci. Each individual is represented by a bar, partitioned into *K* segments representing the extent to which its genome is descended from each ancestral population.

**Table S1.** Isolates and genome sequencing information

**Table S2.** Absolute divergence (dxy) per base pair between *Magnaporthe oryzae* lineages.

**Table S3.** Binary interactions (infection: +; no infection: -; no data: empty cell) between rice varieties from three rice subspecies (left) or rice differential lines (right) and *Magnaporthe oryzae* isolates of six lineages

**Table S4.** General linear model analysis of the proportion of compatible interactions (A) and contrasting results for the analysis of the proportion of compatible interactions (B).

**Table S5.** General linear model analysis of the proportion of R genes overcome (A) and contrasting results for the general linear model analysis of the proportion of R genes overcome (B)

**Table S6.** Distribution of putative migrant mutations, list of predicted genes matching putative migrant mutations and functional enrichment analysis.

